# The chromatin-associated 53BP1 ortholog, HSR-9, regulates recombinational repair and *X* chromosome segregation in the *Caenorhabditis elegans* germ line

**DOI:** 10.1101/2024.04.12.589267

**Authors:** Qianyan Li, Sara Hariri, Aashna Calidas, Arshdeep Kaur, Erica Huey, JoAnne Engebrecht

## Abstract

53BP1 plays a crucial role in regulating DNA damage repair pathway choice and checkpoint signaling in somatic cells; however, its role in meiosis has remained enigmatic. In this study, we demonstrate that the *Caenorhabditis elegans* ortholog of 53BP1, HSR-9, associates with chromatin in both proliferating and meiotic germ cells. Notably, HSR-9 is enriched on the *X* chromosome pair in pachytene oogenic germ cells. HSR-9 is also present at kinetochores during both mitotic and meiotic divisions but does not appear to be essential for monitoring microtubule-kinetochore attachments or tension. Using cytological markers of different steps in recombinational repair, we found that HSR-9 influences the processing of a subset of meiotic double strand breaks into COSA-1-marked crossovers. Additionally, HSR-9 plays a role in meiotic *X* chromosome segregation under conditions where *X* chromosomes fail to pair, synapse, and recombine. Together, these results highlight that chromatin-associated HSR-9 has both conserved and unique functions in the regulation of meiotic chromosome behavior.

**Article Summary:** While 53BP1 is known for its crucial role in DNA damage signaling and repair in somatic cells, its role in meiosis is not well understood. Li, Hariri, et al., show that *C. elegans* 53BP1 not only functions in meiotic recombination and checkpoint signaling but also regulates the transmission of sex chromosomes when meiosis is perturbed. These results highlight the importance of 53BP1 in *C. elegans* meiosis and suggest that 53BP1 has both conserved and organism-specific functions.

## Introduction

The tumor suppressor 53BP1 regulates several biological processes important for genome integrity, including transcription, DNA repair, and chromosome segregation (Mirza-Aghazadeh-Attari et al., 2019). It was originally identified as a p53 interacting protein (<u>53 B</u>inding <u>P</u>rotein <u>1</u>) (Iwabuchi et al., 1994) and was subsequently shown to promote p53-DNA interactions to enhance transcription and checkpoint signaling (Cuella-Martin et al., 2016). Many studies have focused on the role of 53BP1 in DNA repair. 53BP1 binds and protects DNA ends from resection in complex with RIF1, the Shieldin complex, and PTIP, thereby promoting non-homologous end joining (NHEJ), and antagonizes RAD51 filament formation post-resection to inhibit homologous recombination (HR) (Callen et al., 2020; Chapman et al., 2013; Escribano-Diaz & Durocher, 2013; Mirman et al., 2018; Noordermeer et al., 2018; Ward, Minn, van Deursen, et al., 2003). Its role in promoting NHEJ was reinforced by the finding that mutation of 53BP1 can partially suppress the embryonic lethality of *Brca1* mutant mice, which is proposed to be due to the inappropriate use of NHEJ when homologous recombination (HR) is inhibited by *Brca1* mutation (Bouwman et al., 2010; Bunting et al., 2010; Cao et al., 2009; Chen et al., 2020; Li et al., 2016). 53BP1 has also been shown to play important roles in class switch recombination and telomere protection (Difilippantonio et al., 2008; Dimitrova et al., 2008; Rocha et al., 2016; Sundaravinayagam et al., 2019; Ward et al., 2004). In addition, 53BP1 monitors microtubule-kinetochore interactions or tension, which are important for proper chromosome segregation (Jullien et al., 2002; Wang et al., 2017; Yim et al., 2017).

53BP1 is chromatin-associated and becomes enriched at double-stranded breaks (DSBs) together with the histone variant, γ−H2AX (Ward, Minn, Jorda, et al., 2003). In fact, 53BP1 is often used as a marker of DSB formation. At DSBs, 53BP1 has been shown to bind to histone H4 lysine 20 dimethylation (H4K20me2) through its conserved Tudor domain and H2A lysine 15 ubiquitination (H2AK15ub) (Botuyan et al., 2006; Fradet-Turcotte et al., 2013). In yeast, the 53BP1 ortholog, Rad9, associates with histone H3 lysine 79 methylation (H3K79me) at DSBs and regulates strand annealing for crossover recombination (Ferrari et al., 2020). Recent work provides evidence that 53BP1 undergoes liquid-liquid phase separation both in the context of DNA repair and transcriptional regulation (Kilic et al., 2019; Zhang et al., 2022), suggesting that 53BP1 has complex and incompletely understood chromatin interactions.

Recombinational repair in the context of unique chromatin structure is critical for successful meiosis. Despite the known association of 53BP1 with specific chromatin marks (Botuyan et al., 2006; Fradet-Turcotte et al., 2013) and its role in DNA repair (Callen et al., 2020; Chapman et al., 2013; Escribano-Diaz & Durocher, 2013; Mirman et al., 2018; Noordermeer et al., 2018; Ward, Minn, van Deursen, et al., 2003), very few studies have explored the role of 53BP1 in meiosis. Meiotic recombination is initiated by the intentional induction of DSBs by the conserved topoisomerase Spo11 (Dernburg et al., 1998; Keeney et al., 1997). The broken ends are then processed by multiple exonucleases to expose a 3’ single-stranded tail for strand invasion mediated by the recombinases Dmc1 and Rad51 (in many organisms), or RAD-51 alone (in *C. elegans*). Following strand invasion, while both non-crossover and crossover repair outcomes are possible, at least one DSB per chromosome pair must be converted into an interhomolog crossover for proper chromosome segregation (Chen & Weir, 2024; Gartner & Engebrecht, 2022).

53BP1 mutant mice exhibit growth retardation and increased cancer susceptibility yet are fertile, suggesting that 53BP1 does not play an essential role in meiosis (Ward, Minn, van Deursen, et al., 2003). Further, DNA end resection is not affected in mouse spermatocytes lacking 53BP1 (Paiano et al., 2020). In yeast meiosis, Rad9 binds to exogenously induced DSBs but not to Spo11-induced breaks (Usui & Shinohara, 2021). In *C. elegans*, mutation of *hsr-9* (*53BP1*) enhances the phenotype of *brc-1*(*BRCA1*) mutations, instead of suppressing lethality as in mammals, suggesting it may not promote NHEJ (Hariri et al., 2023). HSR-9 has been shown to play a role in repair and signaling of exogenous breaks in the germ line (Ryu et al., 2013); however, its role in meiotic DSB repair or checkpoint signaling has not been reported. Thus, the function of 53BP1 in meiosis remains unclear and may vary across organisms.

Here, we take advantage of the *C. elegans* system to examine the role of HSR-9 in the germ line. We find that HSR-9 is chromatin-associated, enriched on the *X* chromosome pair in oogenic germ cells, and present at kinetochores in cells undergoing mitotic and meiotic divisions. Mutant analyses reveal roles for HSR-9 in the processing of meiotic DSBs into crossovers and in germline apoptosis. Additionally, HSR-9 is important for segregation of the *X* chromosome pair in oogenic germ cells when pairing, synapsis, and recombination are defective.

## Materials and Methods

### Genetics

*C. elegans* strains used in this study are listed in Table S1. Some nematode strains were provided by the Caenorhabditis Genetics Center, which is funded by the National Institutes of Health National Center for Research Resources (NIH NCRR). Strains were maintained at 20°C except for *zyg-1(ts)*, which was maintained at 15°C.

### CRISPR-mediated allele construction

*hsr-9(xoe17)* in *rad-54* and the Hawaiian background used for SNP mapping was generated as previously reported (Hariri et al., 2023). *gfp::v5::hsr-9 (xoe45)* was generated using the CRISPR-Cas9 ribonucleoprotein complexes based on the co-CRISPR method (Paix et al., 2015). A 1324bp double strand DNA fragment was synthesized (Gblock; IDT) and used as the repair template after PCR amplification. This repair template contains sequences for GFP, V5 tag, and flexible linkers, flanked by 5’ and 3’ homology arms of 175 and 198 bps, respectively. Insertion of GFP, V5 and the flexible linkers upstream of the start codon disrupted the PAM sequence, eliminating the need for point mutations to disrupt the guide RNA sequence. Briefly, the Cas9-crRNA-tracrRNA ribonucleoprotein complex, along with the repair template, was microinjected into the *C. elegans* gonad. F1 progenies exhibiting roller/dumpy phenotypes were isolated and genotyped by PCR to confirm GFP insertion. Similarly, *gfp::hsr-9::3xHA (xoe47)* was generated using a 1290bp double-strand DNA repair template to insert the flexible linker-GFP-3xHA sequence before the stop codon. A silent mutation was introduced at the junction of the 5’ homology arm and the flexible linker sequence to disrupt the PAM sequence in the repair template. Guide sequences, repair templates, and genotyping primers for both fusions are provided in Table S2. All strains were outcrossed for a minimum of three times before analyses.

### Embryonic lethality and male self-progeny

L4 stage hermaphrodites were placed on individual plates and allowed to lay eggs. After 24hrs, they were transferred to new plates and this process was repeated for 3 days. Embryonic lethality was determined by counting eggs and hatched larvae 24hrs after removing the adult hermaphrodite and percent was calculated as eggs/(eggs + larvae). Males were scored 72hrs post adult removal and percent was calculated as males/(males + hermaphrodites + eggs).

### Cytological analyses

For live cell imaging, hermaphrodites aged 18–24hrs post L4 stage were anesthetized in 1mM tetramisole (Sigma-Aldrich) and immobilized between a coverslip and a 2% agarose pad on a glass slide. Z-stacks (0.33μm) were captured on a spinning-disk module of the Marianas spinning-disk confocal real-time 3D Confocal-TIRF microscope (Intelligent Imaging Innovations) equipped with an 63x, NA 1.46 objective lens using a Photometrics QuantiEM electron multiplying charge-coupled device (EMCCD) camera, a Zeiss 980 LSM with Airyscan 2 equipped with a 63x 1.4NA objective and the Airyscan Multiplex Super-Resolution-4Y mode, or an API Delta Vision Ultra deconvolution microscope equipped with an 60x, NA 1.49 objective lens. Subsequent Airyscan processing was performed with automatic settings. Z-projections of stacks were generated, cropped, and adjusted for brightness in Fiji.

Immunostaining of germ lines was performed as described (Jaramillo-Lambert et al., 2007) except slides were incubated in 100% ethanol instead of 100% methanol for detection of direct fluorescence of GFP::COSA-1, mCherry::HIM-8, and GFP::RPA-1. For fixed images of GFP::V5::HSR-9, dissected gonads were rapidly immersed in liquid nitrogen and then fixed with 2% paraformaldehyde as in (Janisiw et al., 2020). Primary antibodies are listed in Table S3. Life Technologies secondary donkey anti-rabbit antibodies conjugated to Alexa Fluor 488 or 594 and goat anti-rabbit conjugated to Alexa Fluor 647 were used at 1:500 dilutions. DAPI (2μg/ml; Sigma-Aldrich) was used to counterstain DNA.

Collection of fixed images was performed using an API Delta Vision Ultra deconvolution microscope. Z stacks (0.2 μm) were collected from the entire gonad. A minimum of three germ lines was examined for each condition. Images were deconvolved using Applied Precision SoftWoRx batch deconvolution software and subsequently processed and analyzed using Fiji (ImageJ) (Wayne Rasband, NIH). Images show half-projections of gonads.

To determine the *X*/autosome ratio of GFP::V5::HSR-9 fluorescence, the GFP fluorescence intensity was measured by drawing a region of interest (ROI) around the *X* chromosomes in pachytene nuclei. The intensity was also measured using the same ROI in an area within the same nucleus that excluded the *X* chromosomes. For H3S10P quantification, nuclei positive for H3S10P signal were counted in the proliferative zone of germ lines from age-matched 18hr post-L4 hermaphrodites grown at 25C to inactivate *zyg-1*. GFP::RPA-1 fluorescence was quantified by measuring the mean fluorescence intensity and standard deviation (SD) in Fiji for individual nuclei from transition zone to mid-pachytene. Coefficient of variation (CV) was calculated as SD of intensity divided by mean intensity (Bishop et al., 2015). For RAD-51 quantification, germ lines were divided into the transition zone (leptotene/zygotene, from the first to last row with two or more crescent-shaped nuclei), and pachytene (divided into 3 equal parts: early, mid, and late pachytene). RAD-51 foci per nucleus was scored from half projections of the germ lines for each divided region. CHK-1Ser345p foci were quantified in early pachytene nuclei. GFP::COSA-1 foci were scored from deconvolved 3D z-stacks in mid-late pachytene nuclei individually to ensure that all foci within each individual nucleus were counted. mCherry::HIM-8 foci were quantified from deconvolved 3D z-stacks throughout the germline.

### Meiotic mapping

Meiotic crossover frequencies and distribution were assayed utilizing single-nucleotide polymorphism (SNP) markers as in (Nabeshima et al., 2004). The SNP markers located at the boundaries of the chromosome domains were chosen based on data from WormBase (WS231), (Bazan & Hillers, 2011), and (Saito et al., 2013) and are indicated in Figure 5D. The SNP markers and primers used are listed in (Li et al., 2020). PCR and restriction digests of single worm lysates were performed as described in (Li et al., 2020).

### X chromosome nondisjunction

Hybrid hermaphrodites expressing GFP::2xNLS and tdTomato::H2B (fusions inserted into the same location on each of the *X* chromosomes) were mated to *fem-3(e1996)/*nT1GFP; *lon-2(e678)* males. Parents were transferred every 24hrs for 3 days. Progeny were scored for GFP and tdTomato on a fluorescent stereo microscope at 20x magnification on day 3. Only worms with green-fluorescent pharynxes were scored for *X* chromosome markers to ensure cross progeny were examined.

### Statistical analyses

Statistical analyses and figures were prepared using GraphPad Prism version 10.0 (GraphPad Software). Statistical comparisons of embryonic lethality (Figure S1B, Figure S2A, and Figure 6A), *X*/autosomal GFP::V5::HSR-9 fluorescence intensity (Figure 2E), H3S10P nuclei/germ line (Figure 3D), GFP::RPA-1 fluorescence (Figure 4C and Figure S2B), RAD-51 foci numbers (Figure 4E, G and Figure S2C), % male self-progeny, pCHK-1 foci/nucleus and apoptotic nuclei/gonad (Figure 6B-E) were analyzed by Mann-Whitney. Chi-squared test was used for statistical analyses on GFP::COSA-1, genetic map distance, distribution, and % oocytes with X chromosome non-disjunction events (Figure 5A, D, and Figure 7B). Detailed descriptions of statistical analyses are indicated in figure legends.

**Figure 1.**
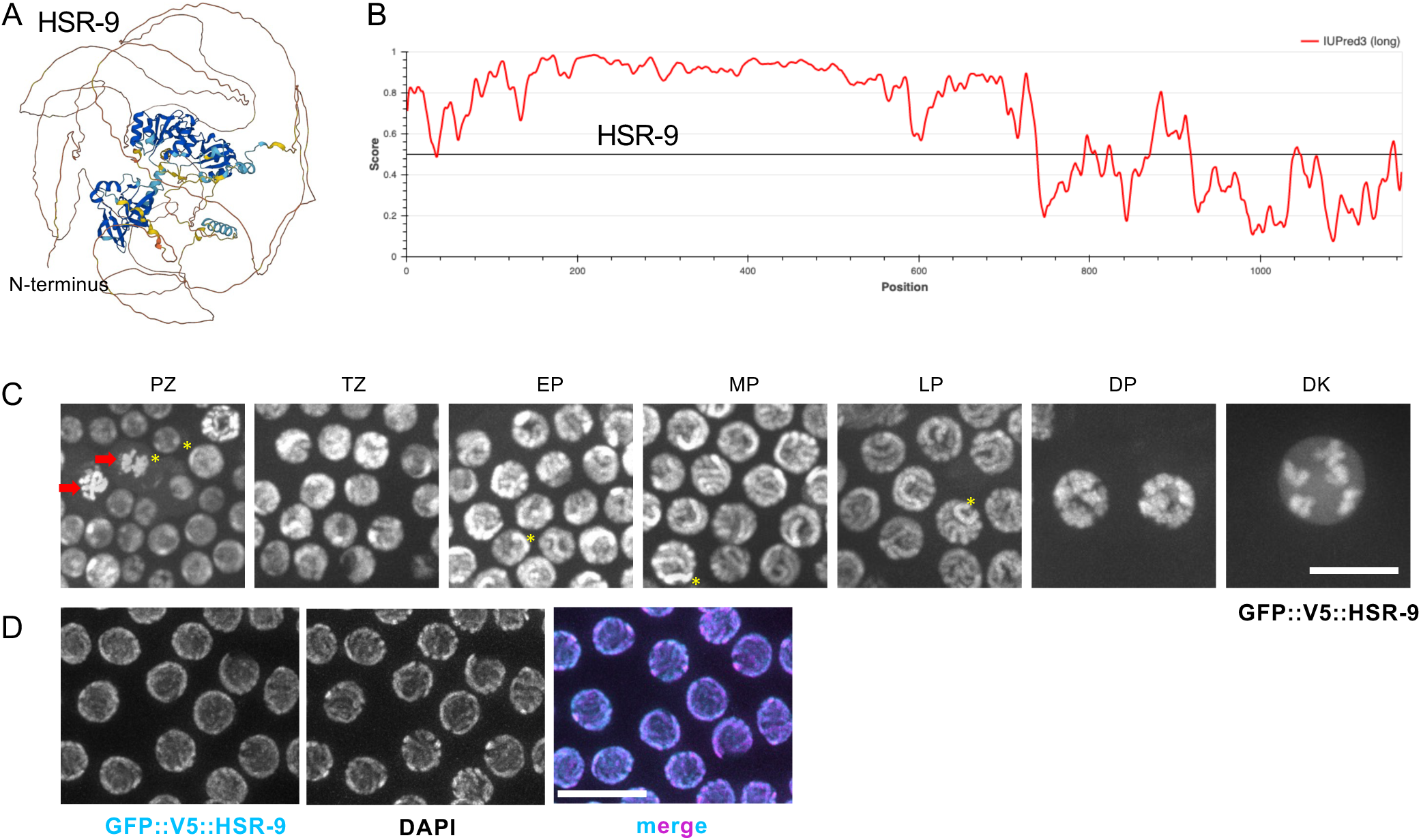
HSR-9 is a chromatin-associated protein with intrinsically disordered N-terminal region. A) Predicted structure of HSR-9 determined by AlphaFold (Jumper et al., 2021). N-terminus is indicated in brown line. B) Predicted disordered regions of HSR-9 using IUPred3 (Erdos et al., 2021). C) Images of GFP::V5::HSR-9 fluorescence from indicated regions of the germ line. Red arrows mark condensing chromosomes in prometaphase to anaphase, asterisks mark two domains with stronger fluorescense intensity. PZ = proliferative zone; TZ = transition zone; EP = early pachytene; MP = mid pachytene; LP = late pachytene; DP = diplotene; DK = diakinesis. Scale bar = 10μm. D) Images of GFP::V5::HSR-9 fluorescence of the pachytene region of dissected gonads following rapid freeze crack, fixation in cold ethanol and paraformaldehyde treatment counterstained with DAPI. Scale bar = 10μm.

**Figure 2.**
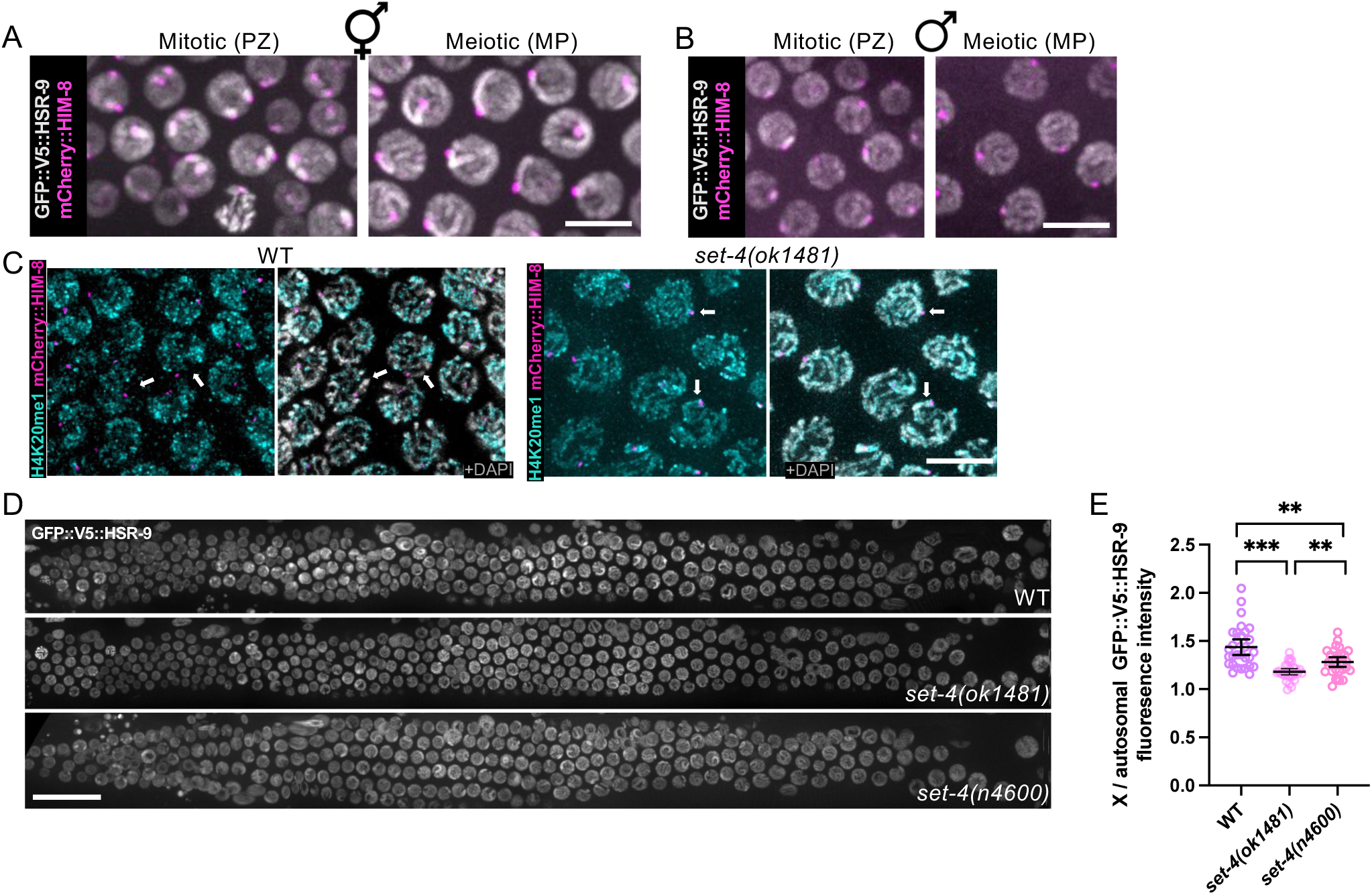
GFP::V5::HSR-9 is enriched on *X* chromosomes in oogenesis and associates with H4K20me2/me3. Images of GFP::V5::HSR-9 (grey) and mCherry::HIM-8 (magenta) from mitotic (PZ, proliferative zone) and meiotic (MP, mid-pachytene) regions of the germ line in live A) hermaphrodites and B) males. C) Images of fixed and dissected meiotic germ cells labelled with antibodies against H4K20me1 (cyan) in wild type and *set-4(ok1481)* expressing mCherry::HIM-8 (magenta) and counter-stained with DAPI (grey). Arrows point to the X chromosomes. Scale bar = 5μm. D) GFP::V5::HSR-9 fluorescence of whole gonads from live worms from wild type, *set-4(ok1481)*, and *set-4(n4600).* Scale bar = 20 μm. E) Quantification of the relative fluorescence intensity of the *X* to autosome ratio (n=3 worms for each genotype). *** p < 0.001; ** p < 0.01 by Mann-Whitney.

**Figure 3.**
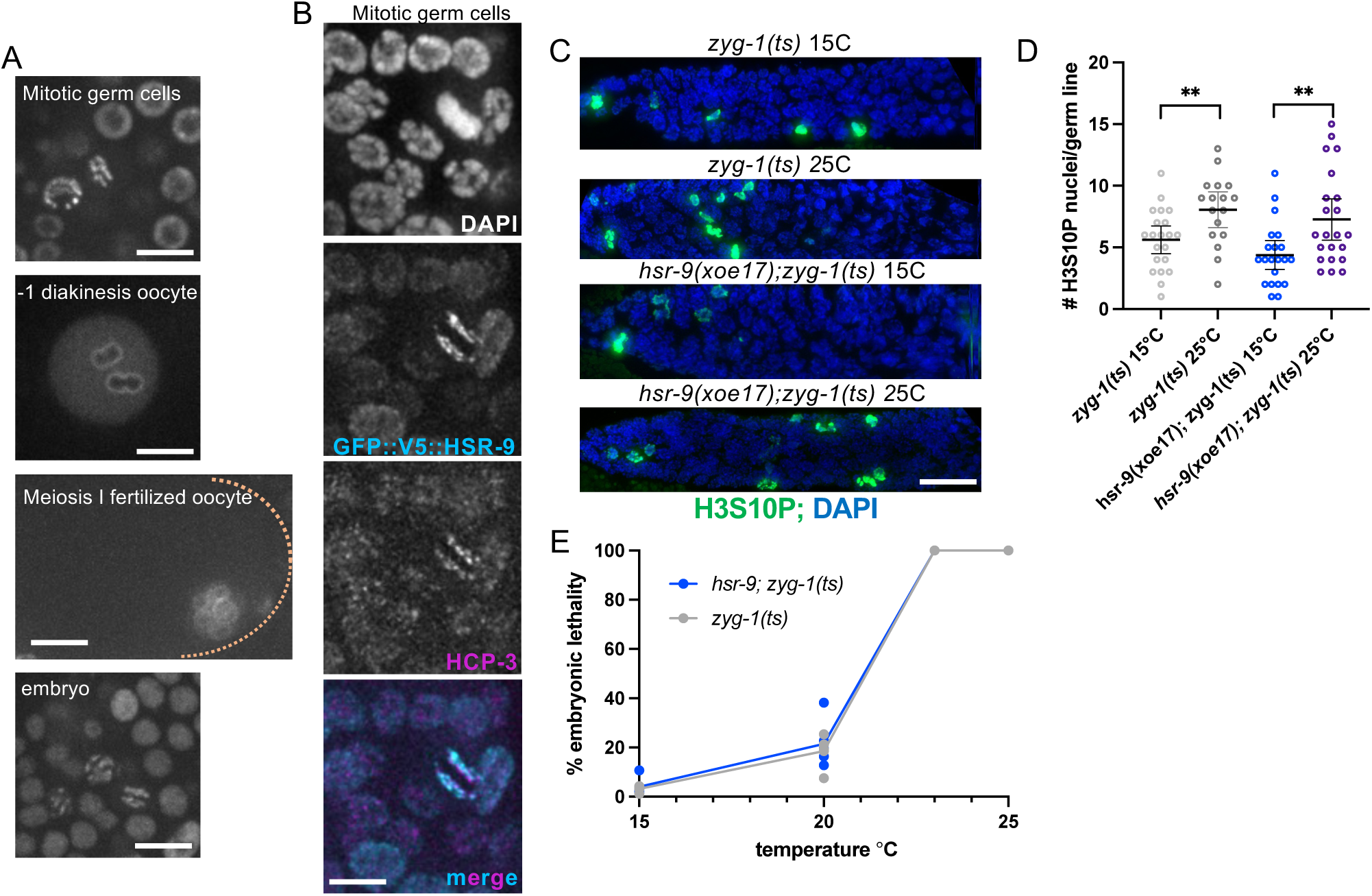
GFP::V5::HSR-9 is enriched on kinetochores but doesn’t play an essential role in monitoring microtubule-kinetochore attachments. A) Live imaging of GFP::V5::HSR-9 fluorescence in mitotic germ cells, −1 diakinesis oocyte, Meiosis I fertilized oocyte and embryo showing structures consistent with enrichment on kinetochores. B) Fixed images of mitotic germ cells labelled with CENPA ortholog, HCP-3 (magenta) and direct GFP fluorescence (cyan). Scale bar = 5μm. C) Fixed images of mitotic region of germ line labelled with antibodies directed against H3S10P and counterstained with DAPI at indicated temperatures. Scale bar = 20 μm. D) Quantification of H3S10P nuclei/germ line in *zyg-1(ts)* and *hsr-9(xoe17); zyg-1(ts)* grown at 15° or 25°C. Number of germ lines examined: *zyg-1(ts)* 15°C = 21; *zyg-1(ts)* 25°C = 21; hsr-9(xoe17); *zyg-1(ts)* 15°C = 21; *hsr-9(xoe17); zyg-1(ts)* 15°C = 22; ** p < 0.01; by Mann-Whitney. E) % embryonic lethality of *zyg-1(ts)* and *hsr-9(xoe17); zyg-1(ts)*; 6 worms were examined at each temperature.

**Figure 4.**
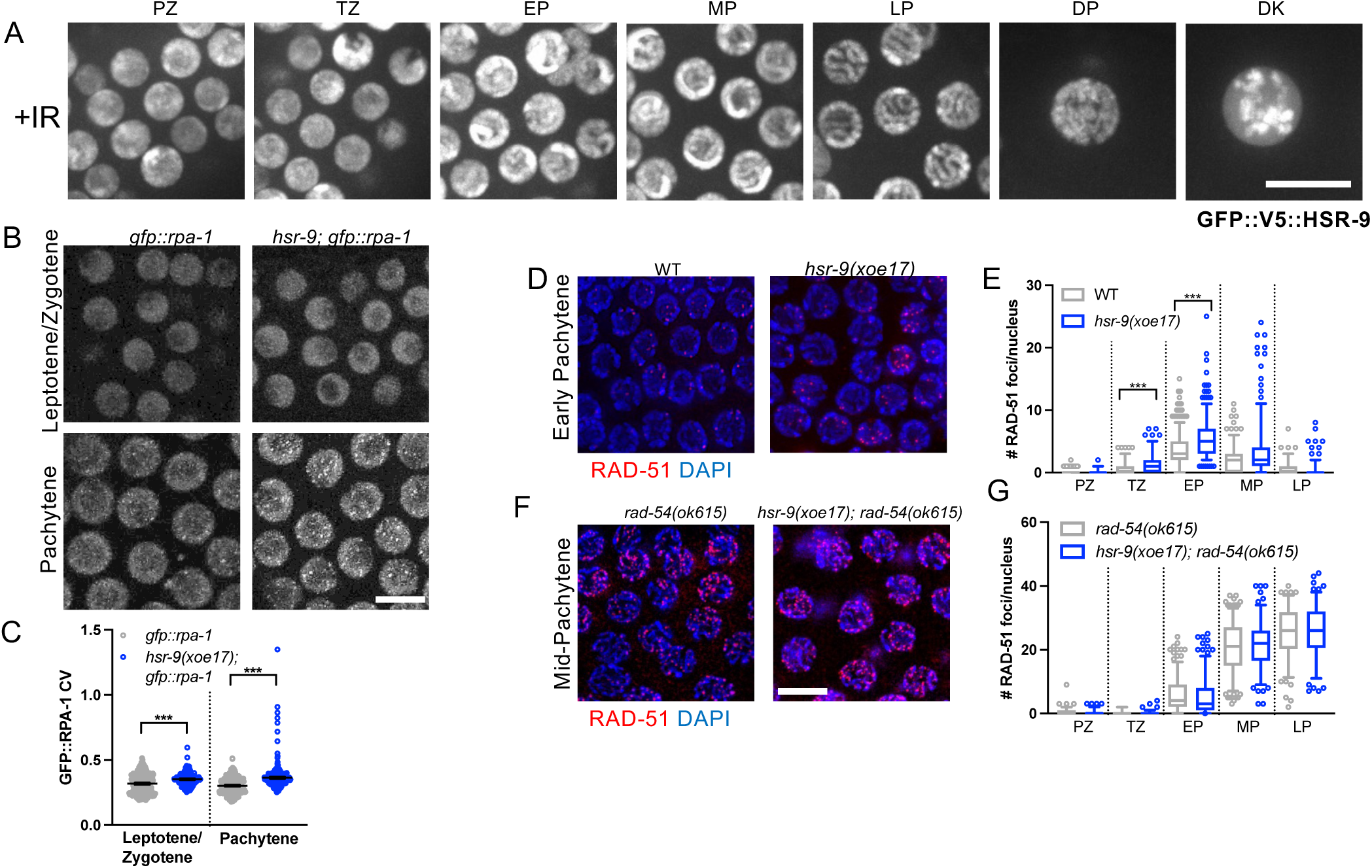
HSR-9 plays a role in processing of meiotic DSBs. A) Images of GFP::V5::HSR-9 fluorescence from indicated regions of the germ line 1hr post 75Gys IR. PZ = proliferative zone; TZ = transition zone; EP = early pachytene; MP = mid pachytene; LP = late pachytene; DP = diplotene; DK = diakinesis. Scale bar = 10μm. B) GFP-RPA-1 fluorescence in leptotene/zygotene and pachytene. Scale bar = 5μm. C) Coefficient of Variation (CV) of GFP::RPA-1 fluorescence was measured from 3 germ lines. Images of early pachytene nuclei immunolabelled with RAD-51 (red) and counter stained with DAPI (blue) from wild type (WT) and *hsr-9(xoe17)* germ lines (D) and corresponding quantification (E), and *rad-54(ok615)* and *hsr-9(xoe17); rad-54(ok615)* (F) and corresponding quantification (G). Box whisker plots show number of RAD-51 foci per nucleus in the indicated regions. Horizontal line of each box represents the median, top and bottom of each box represent medians of upper and lower quartiles, lines extending above and below boxes indicate SD, and individual data points are outliers from 5 to 95%. Statistically significant comparisons by Mann–Whitney of WT vs. *hsr-9(xoe17)* are indicated; ***p< 0.0001. Complete germlines are shown in Figure S2.

**Figure 5.**
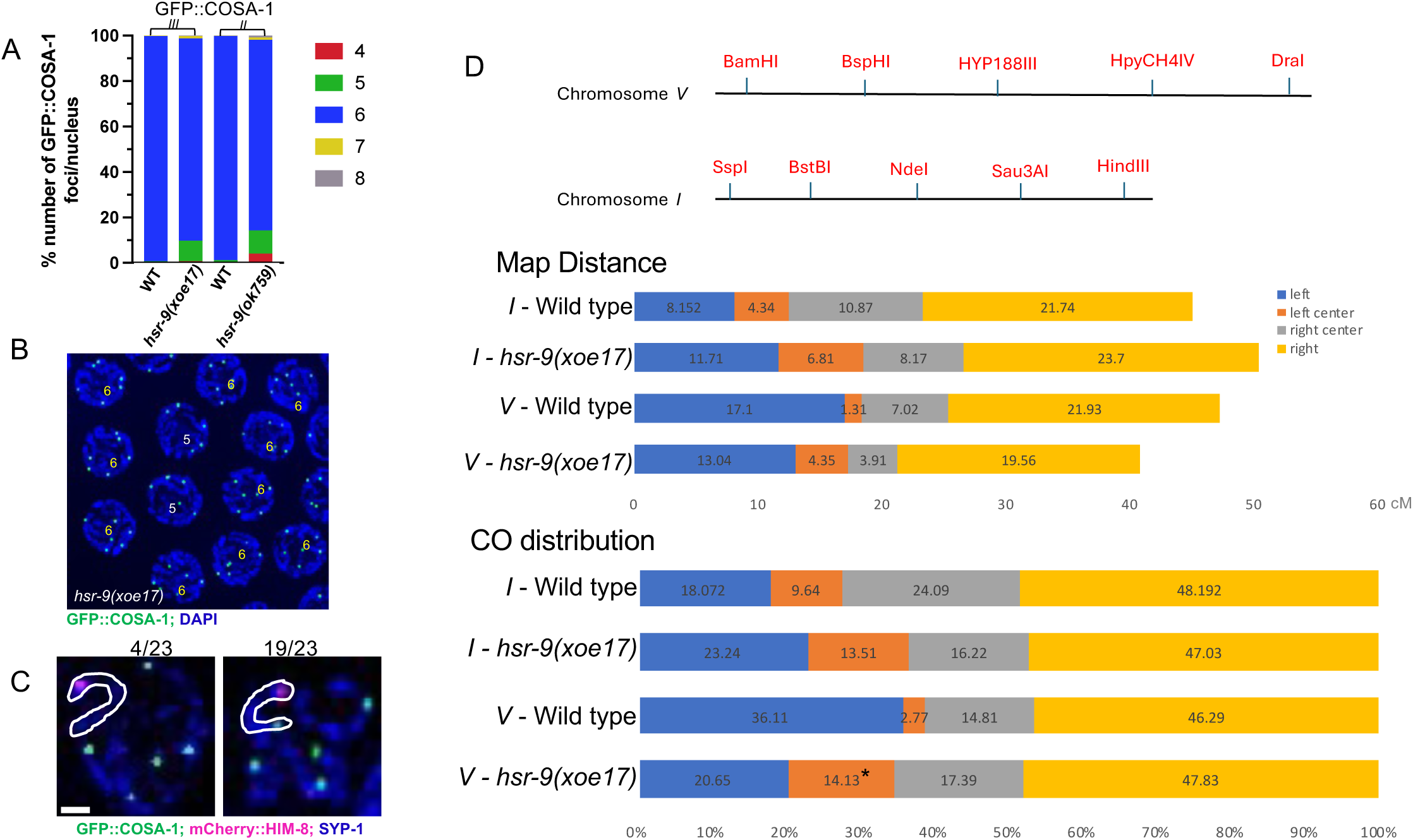
Reduced numbers of GFP::COSA-1, but not genetic crossovers in the absence of HSR-9. A) Percent of nuclei containing indicated GFP::COSA-1 foci in *gfp::cosa-1(xoe44)* and *hsr-9(xoe17); gfp::cosa-1(xoe44),* which is at the endogenous locus on chromosome *III* and *meIs8[unc-119(+) pie-1promoter::gfp::cosa-1]* and *hsr-9(ok759); meIs8[unc-119(+) pie-1promoter::gfp::cosa-1]* inserted on chromosome *II*. B) Image of mid-late pachytene nuclei showing GFP::COSA-1(green) and counterstained with DAPI (blue). Number of GFP::COSA-1 foci are indicated on each nucleus. Scale bar = 5μm. C) Images of pachytene nuclei immunolabelled with SYP-1 antibodies (blue), and imaged for mCherry::HIM-8 (magenta) and GFP::COSA-1 (green) fluorescence. Scale bar = 1 μm. D) Top: SNP markers (red) on Chromosome *I* and *V*. Middle: crossover frequency on Chromosomes *I* and *V*. Bottom: crossover distribution among recombinants on Chromosomes *I* and *V*. Total number of worms analyzed for Chromosome *I* markers = wild type (n = 184), and *hsr-9(xoe17)* (n = 367); Chromosome *V* markers = wild type (n = 228), and *hsr-9(xoe17)* (n = 230). Statistical analyses were conducted using χ2 on 2-by-2 contingency tables,* p<0.05.

**Figure 6.**
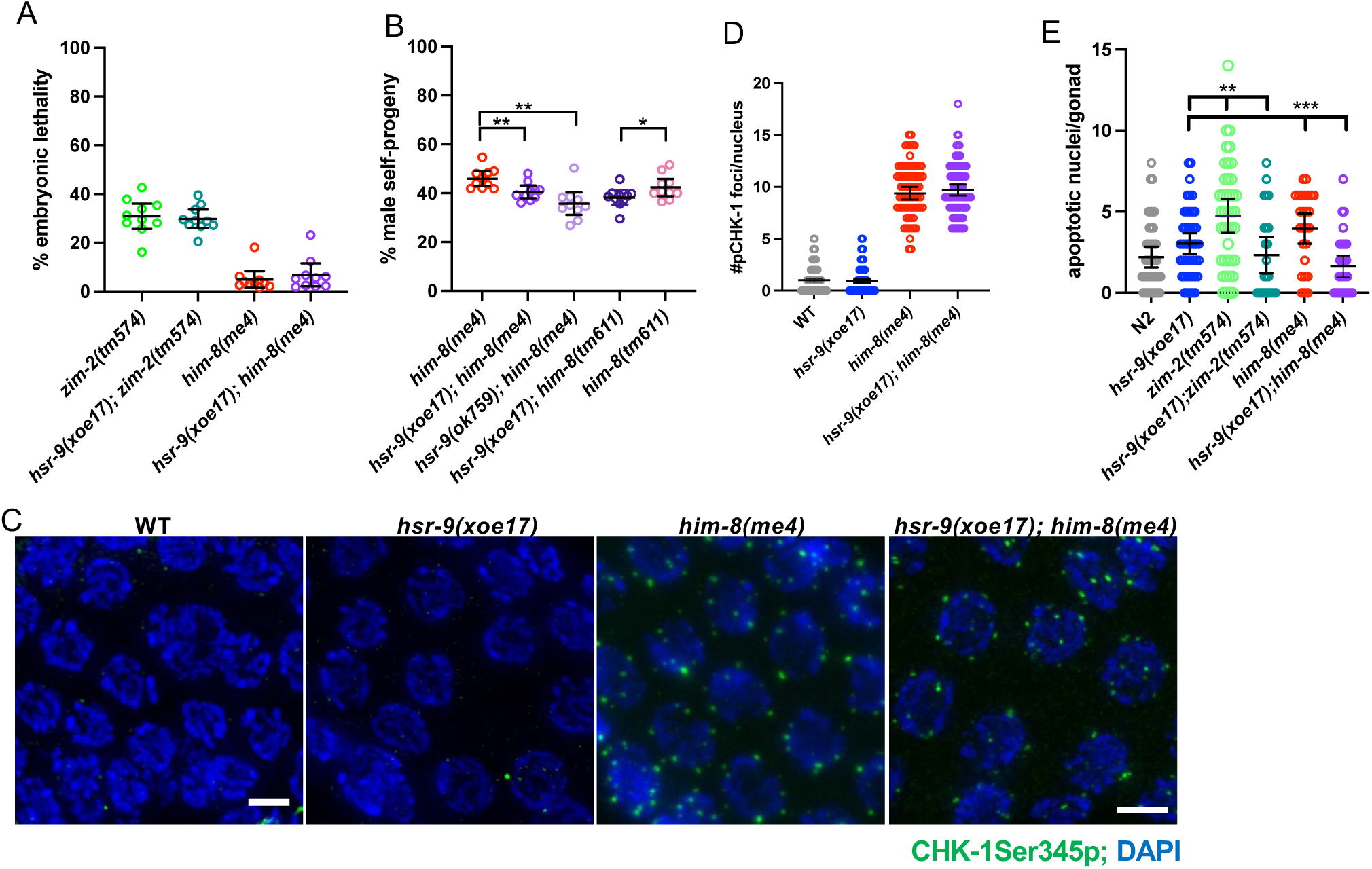
Apoptosis is reduced but checkpoint signaling appears intact in *hsr-9* mutants. A) Embryonic lethality in *zim-2(tm574)*, *hsr-9(xoe17); zim-2(tm574)*, *him-8(me4)* and *hsr-9(xoe17); him-8(me4)* mutants (n=10). B) % male self-progeny in him-*8(me4), hsr-9(xoe17); him-8(me4), hsr-9(ok759); him-8(me4), hsr-9(xoe17); him-8(tm611)* and *him-8(tm611)* mutants (n=10). C) Early pachytene germ cells immunolabelled with CHK-1Ser345p (green) in WT, *hsr-9(xoe17)*, *him-8(me4)* and *hsr-9(xoe17); him-8(me4)*. Scale bar = 5μm. D) Number of CHK-1Ser345p foci per nuclei in WT (n=187), *hsr-9(xoe17)* (n=200), *him-8(me4)* (n=75) and *hsr-9(xoe17); him-8(me4)* n=87). E) Number of apoptotic nuclei/gonad by Acridine Orange staining in WT (n=44), *hsr-9(xoe17)* (n=45) *zim-2(tm574)* (n=41), *hsr-9(xoe17); zim-2(tm574)* (n=24), *him-8(me4)* (n=26) and *hsr-9(xoe17); him-8(me4)* (n=32). * p<0.05; ** p<0.001; *** p<0.0001 by Mann-Whitney.

**Figure 7.**
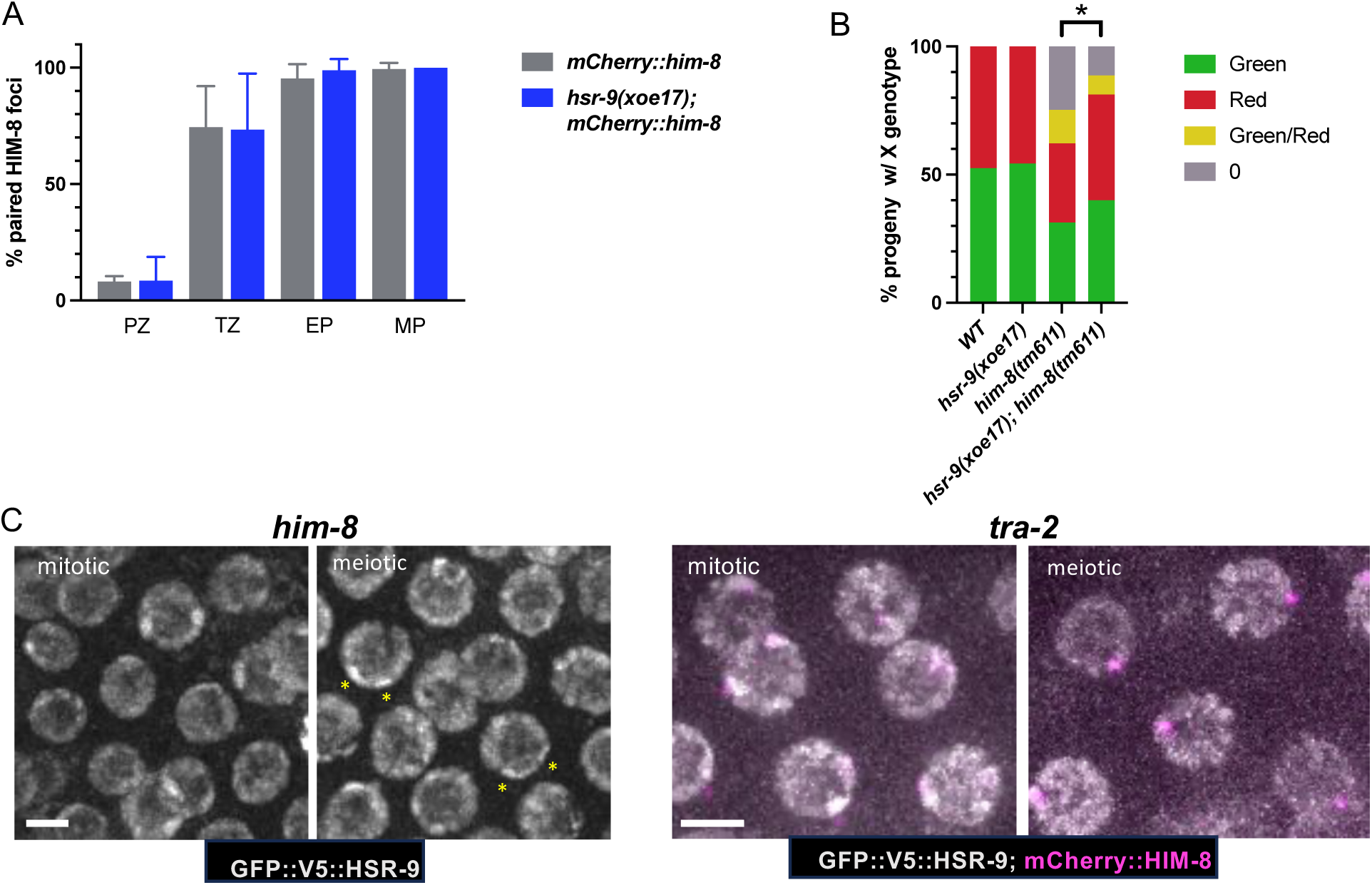
HSR-9 plays a role in *X* chromosome segregation in *him-8*. A) % nuclei with paired (1 focus) mCherry::HIM-8 foci in *mCherry::him-8* and *hsr-9(xoe17); mCherry::him-8* worms at the indicated stages of meiotic prophase: PZ, proliferative zone, TZ, transition zone; EP, early pachytene; MP, mid pachytene. Three gonads were analyzed for each genotype. B) % oocytes with *X* chromosome expressing GFP::2xNLS (green), tdTomato::H2B (red), both GFP::2xNLS and tdTomato::H2B (*XX* - yellow) or no *X* chromosome (nullo-*X* - grey). Number of progeny examined: WT = 148, *hsr-9(xoe17)* = 196, *him-8(tm611)* = 145, *hsr-9(xoe17); him-8(tm611)* = 163. * p<0.05 by χ2 on 2-by-4 contingency tables C) Images of GFP::V5::HSR-9 (grey) in *him-8* (yellow asterisks denote regions of more intense fluorescence) and GFP::V5::HSR-9 (grey) and mCherry::HIM-8 (magenta) in *tra-2 XX* males from mitotic and meiotic regions of the germ line in live worms. Scale bar = 5μm.

## Results

### HSR-9 associates with chromatin in the germ line

HSR-9 exhibits a domain structure similar to human 53BP1, with an intrinsically disordered N-terminus and conserved BRCT domains at the C-terminus. The predicted structure of HSR-9 as determined by AlphaFold is shown in Figure 1A (Jumper et al., 2021) and the extent of disorder is shown in Figure 1B (Erdos et al., 2021). To examine HSR-9 localization, we constructed GFP fusions to both the N and C-termini of HSR-9 using CRISPR genome editing (Figure S1A). Mutation of HSR-9 has no effect on progeny viability (Ryu et al., 2013), and we confirmed this using the putative null allele *hsr-9(xoe17)* (Hariri et al., 2023) (Figure S1A). We examined the functionality of the GFP fusions to HSR-9 by monitoring embryonic lethality in the absence of the ortholog of the tumor suppressor BRCA1/BRC-1, as the *hsr-9; brc-1* double mutant results in enhanced embryonic lethality [(Hariri et al., 2023); Figure S1B]. The C-terminal HSR-9::GFP::3xHA fusion in combination with *brc-1* showed similar embryonic lethality to the *hsr-9; brc-1* double mutant, suggesting it is nonfunctional. On the other hand, *gfp::v5::hsr-9; brc-1* double mutant exhibited embryonic lethality at levels between that of wild type and *hsr-9; brc-1,* suggesting that GFP::V5::HSR-9 is partially functional (Figure S1C). Both fusions showed similar localization in the *C. elegans* germ line (Figure 1C and Figure S1C). We subsequently concentrated our localization studies on worms expressing the GFP::V5::HSR-9 fusion.

We monitored HSR-9 localization by live imaging focusing on the *C. elegans* germ line, which contains proliferating germ cells at the distal end and all stages of meiotic prophase arranged in a spatial-temporal gradient towards the proximal end (Figure 1C). Consistent with a previous study using an antibody directed against HSR-9 (Ryu et al., 2013), GFP fluorescence was enriched in the nucleus of proliferating germ cells (Figure 1C). In metaphase and anaphase of mitosis, where the nuclear envelope breaks down and chromosomes condense, HSR-9 was localized to the condensed chromosomes (PZ, arrows; Figure 1C), suggesting that the protein is chromatin-associated. In meiotic germ cells, HSR-9 was observed on chromatin throughout prophase, first appearing in tracks as chromosomes synapse, and subsequently on condensed chromosomes in diplotene/diakinesis (Figure 1C).

In contrast to observations from live imaging, the GFP::V5::HSR-9 signal in dissected and fixed gonads did not show as tight an association with chromatin (Ryu et al., 2013). This pattern held true regardless of whether we used antibodies directed against GFP or V5, or imaged GFP fluorescence directly on samples prepared by dissection in paraformaldehyde followed by ethanol fixation. Rapid freeze crack and fixation in cold ethanol followed by paraformaldehyde treatment of dissected gonads more closely but not entirely recapitulated the results by live imaging (Figure 1D). As proteins containing intrinsically disordered regions have been shown to have different localization patterns in fixed versus live cells, the difference we observed for HSR-9 localization is likely a consequence of rapid association/disassociation of HSR-9 with chromatin (Irgen-Gioro et al., 2022; Schmiedeberg et al., 2009; Teves et al., 2016).

### HSR-9 is enriched on the *X* chromosome pair in hermaphrodite germ cells

We noted that GFP::V5::HSR-9 fluorescence in live hermaphrodite worms was more intense in two nuclear domains in many mitotic germ cells where homologous chromosomes are unpaired, and on one chromosome track in pachytene germ cells where homologous chromosomes are paired and synapsed (asterisks, Figure 1C). We hypothesized that the GFP::V5::HSR-9-enriched regions were the *X* chromosomes, which have different chromatin properties compared to the autosomes (Bean et al., 2004; Checchi & Engebrecht, 2011; Jaramillo-Lambert & Engebrecht, 2010; Kelly et al., 2002; Reuben & Lin, 2002). To test this, we imaged live worms expressing mCherry::HIM-8, the *X* chromosome-specific pairing center protein (Link et al., 2018; Phillips et al., 2005), and found that mCherry::HIM-8 labelled the two GFP::V5::HSR-9 enriched domains in many proliferating germ cells as well as the single chromosome track enriched for GFP::V5::HSR-9 in pachytene germ cells (Figure 2A). This result is consistent with GFP::V5::HSR-9 being concentrated on *X* chromosomes. Interestingly, while we observed a single HIM-8-associated GFP::V5::HSR-9 enriched region in mitotic germ cells in the male germ line, consistent with enrichment on the single *X* chromosome of males, we did not observe enrichment on the *X* chromosome in male pachytene germ cells (Figure 2B). Thus, GFP::V5::HSR-9 is associated with chromatin, and is enriched on the *X* chromosome(s) in both oogenic and spermatogenic proliferating germ cells but only on the *X* chromosome pair in oogenic pachytene germ cells.

Mammalian 53BP1 associates with histone H4 dimethylated on lysine 20 (H4K20me2). In *C. elegans* pachytene germ cells, H4K20me1 is enriched on autosomes relative to the *X* chromosomes due to autosomal targeting of the DPY-21 demethylase, which converts H4K20me2 to H4K20me1 (Brejc et al., 2017; Vielle et al., 2012). This suggests that *X* chromosomes are enriched for H4K20me2/3 and this enrichment may contribute to HSR-9 accumulation on the *X* chromosomes. As we were unable to identify antibodies specific for H4K20me2/3, we labelled germ lines with antibodies against H4K20me1 and observed enrichment on autosomes relative to the *X* chromosomes as previously reported (Brejc et al., 2017; Vielle et al., 2012) (Figure 2C). We next examined the immunolocalization of H4K20me1 in two different deletion mutants of the SET-4 methyltransferase, which adds methyl groups to H4K20me1 to generate H4K20me2 and H4K20me3 (Kramer et al., 2015; Vielle et al., 2012; Wells et al., 2012). *set-4(ok1481)* contains a 913bp C-terminal deletion that removes that last 17 amino acids and 3’UTR, while *set-4(n4600*) contains a 1146bp deletion that removes upstream sequences and the first 165 of 288 amino acids. In both mutants, H4K20me1 was detectable on the *X* chromosome pair, consistent with the *X* chromosomes being enriched for SET-4-dependent H4K20me2/me3 in wild-type animals (Figure 2C). Analysis of GFP::V5::HSR-9 in *set-4* mutants revealed a reduced, but still detectable, enrichment of GFP::V5::HSR-9 on the *X* chromosome pair (Figure 2D, E). These results suggest that H4K20me2/me3 contributes to the enrichment of HSR-9 on the *X* chromosome pair, but there are additional factors, likely including other chromatin marks, that influence HSR-9 recruitment.

### HSR-9 localizes to kinetochores but does not play an essential role in monitoring kinetochore-spindle attachments/tension

In mammalian somatic cells, 53BP1 accumulates at kinetochores and centrosomes in addition to being associated with chromatin (Jullien et al., 2002; Yim et al., 2017). *C. elegans* chromosomes are holocentric, with kinetochore proteins binding along the poleward face of each sister chromatid during mitosis and forming cup-shaped structures on meiotic chromosomes (Albertson & Thomson, 1982; Monen et al., 2005). While no centrosome localization was detected, we observed GFP::V5::HSR-9 in bar-like structures on the poleward face of metaphase chromosomes in proliferating germ cells and in embryos (Figure 3A). To confirm that GFP::V5::HSR-9 is enriched on kinetochores, we performed immunolabeling with antibodies against the kinetochore-associated CENP-A ortholog, HCP-3 (Gassmann et al., 2012) and imaged GFP fluorescence. We observed both HCP-3 and GFP fluorescence in bar-like structures that co-localized on the poleward face of metaphase chromosomes in proliferating germ cells (Figure 3B), confirming that HSR-9 is enriched at kinetochores. GFP::V5::HSR-9 was also observed surrounding the outer edge of chromosomes at the −1 oocyte and at the meiotic divisions (Figure 3A), consistent with kinetochore localization.

Mammalian 53BP1 at the kinetochore has been shown to play a role in monitoring inappropriate microtubule attachments or tension (Jullien et al., 2002; Wang et al., 2017). To determine whether HSR-9 plays a similar role in germ cells, we disrupted metaphase using the *zyg-1(b1)* conditional mutant [referred to as *zyg-1(ts)*; (O’Connell et al., 2001; Wood et al., 1980)], which we previously showed perturbed spindle function in mitotically-dividing germ cells and activated the DNA damage response and spindle assembly checkpoint (Lawrence et al., 2015). ZYG-1 is functionally related to PLK4 and is required for centrosome duplication (O’Connell et al., 2001). Its inactivation leads to the formation of monopolar spindles and disrupts spindle attachments and tension. This results in a cell cycle delay, which is evidenced by an increase in the number of nuclei enriched for phosphorylation of Serine 10 on Histone H3 (H3S10P), a marker of prometaphase/metaphase (Lawrence et al., 2015; Prigent & Dimitrov, 2003). As expected, we observed an increase in H3S10P-positive nuclei following ZYG-1 inactivation at 25°C, indicative of a metaphase delay (Figure 3C, D). Inactivation of *zyg-1(ts)* at the nonpermissive temperature in the *hsr-9(xoe17)* mutant did not alter the number of H3S10P-positive nuclei, suggesting that HSR-9 is not required for monitoring microtubule attachment/tension and checkpoint activation in the *zyg-1(ts)* mutant (Figure 3C, D). We also examined the consequence of HSR-9 mutation on progeny viability at different temperatures in the *zyg-1(ts)* mutant. No enhancement of progeny lethality was observed in the absence of HSR-9, consistent with our findings that HSR-9 is not essential for cell cycle delay when monopolar spindles are induced. Thus, HSR-9 is enriched at kinetochores in metaphase but does not play a significant role in monitoring spindle attachments/tension at kinetochores in *C. elegans*.

### HSR-9 regulates meiotic DSB processing

In somatic cells, 53BP1 functions in the DNA damage response and repair choice (Mirman & de Lange, 2020). Further, 53BP1 becomes enriched in nuclear foci following DNA damage, and has been widely used as a marker of DSBs (Ward, Minn, Jorda, et al., 2003). No obvious GFP::V5::HSR-9 foci were observed in meiotic cells, where DSBs are induced by SPO-11 (Figure 1C). Further, no GFP::V5::HSR-9 foci were observed following irradiation (IR) treatment, suggesting that HSR-9 does not accumulate at DSBs in *C. elegans* germ cells (Ryu et al., 2013) (Figure 4A).

While HSR-9 does not accumulate at DSBs, HSR-9 has been reported to play a role in repair of IR-induced breaks in the germ line when HR is impaired (Ryu et al., 2013). To determine whether HSR-9 functions in repair of meiotic DSBs, we analyzed meiotic recombination in *hsr-9(xoe17)* and the previously described *hsr-9(ok759)* allele, which removes 1613 bps in the middle of the gene (Ryu et al., 2013) (Figure S1A). Progeny viability was high (Figure S1B) and there was no increase in male self-progeny in either *hsr-9(ok759)* or *hsr-9(xoe17)* mutants (wt = 0.02±0.01, *hsr-9(ok759)* = 0.01±0.05, *hsr-9(xoe17)* = 0.01±0.04% males), suggesting that HSR-9 is not essential for meiotic recombination. Further, blocking apoptosis by mutation of CED-3, the caspase essential for executing cell death (Ellis & Horvitz, 1986; Yuan et al., 1993), did not increase the number of inviable progeny compared to the *ced-3* mutant alone, suggesting that the lack of elevated progeny lethality is not due to culling by apoptosis in the absence of HSR-9 (Figure S2A).

Meiotic DSBs are catalyzed by the conserved topoisomerase SPO-11 (Bergerat et al., 1997; Dernburg et al., 1998; Keeney et al., 1997), and then processed for repair predominately by HR. We monitored meiotic DSB repair by examining the appearance and disappearance of the Replication Protein A (RPA) complex and RAD-51 in the spatiotemporal organization of the *C. elegans* germ line. RPA coats single-stranded DNA generated by end resection. In *C. elegans*, RPA is composed of RPA-1 and RPA-2 (Hefel et al., 2021) and GFP::RPA-1 is observed in foci from early prophase (leptotene/zygotene) through pachytene, suggesting that it not only marks resected ends but also recombination intermediates post-resection (Woglar & Villeneuve, 2018). In live worms, GFP::RPA-1 is nucleoplasmic with some nuclear foci visible (Li et al., 2020; Li et al., 2023). Live cell imaging of GFP::RPA-1 in wild type and *hsr-9(xoe17)* revealed more intense foci in the *hsr-9(xoe17)* mutant (Figure 4B). To quantify this, we measured the coefficient of variation (CV), which describes the dispersion of pixel intensity values from a 2D region of interest around the mean pixel intensity such that nuclei with more foci above the nucleoplasmic signal will have high CV values, whereas nuclei with more uniform fluorescence will have low CV values (Bishop et al., 2015). We observed a higher CV of GFP::RPA-1 in *hsr-9(xoe17)* compared to wild type in both leptotene/zygotene and pachytene nuclei (Figure 4C). GFP::RPA-1 fluorescence also had a higher CV in the *hsr-9(ok759)* mutant (Figure S2B).

We next examined the assembly and disassembly of RAD-51 (Rinaldo et al., 2002) in the presence and absence of HSR-9 using antibodies against RAD-51 (Alpi et al., 2003; Colaiacovo et al., 2003). RAD-51 replaces the RPA complex on resected DSBs beginning in leptotene/zygotene (transition zone) and is largely removed by late pachytene (Colaiacovo et al., 2003). While the overall pattern of RAD-51 is similar in *hsr-9* mutant vs. wild-type germ lines, there was a significant increase in the number of RAD-51 foci detected in early meiotic prophase (transition zone and early pachytene) in both *hsr-9(xoe17)* and *hsr-9(ok759)* mutants (Figure 4D, E; Figure S2C). Elevated RPA-1 and RAD-51 foci could be a consequence of a greater number of DSBs repaired by HR, and/or a defect in processing of DSBs.

To provide insight into the nature of the elevated RAD-51 foci in *hsr-9* mutants, we analyzed RAD-51 in the absence of RAD-54. *rad-54* mutants have been used to distinguish between increased number of DSBs vs. a defect in processing of breaks as RAD-54 is essential for RAD-51-mediated strand exchange during HR and is required for RAD-51 disassembly; in its absence RAD-51 remains on processed breaks (Mets & Meyer, 2009; Solinger et al., 2002). The number of RAD-51 foci per nucleus in *rad-54* mutants in the presence or absence of HSR-9 was not statistically different (Figure 4F, G and Figure S2D), suggesting that HSR-9 does not alter the number of DSBs formed but rather regulates the processing of DSBs by HR.

### Not all crossovers accumulate COSA-1 in the absence of HSR-9

To determine whether the defect in processing of DSBs alters crossovers in the *hsr-9* mutants, we monitored the <u>c</u>ross<u>o</u>ver <u>s</u>ite <u>a</u>ssociated protein COSA-1/CNTD1 (Yokoo et al., 2012). Wild-type hermaphrodites have six GFP::COSA-1 foci per nucleus, one on each of the six pairs of homologous chromosomes, in mid-late pachytene. In *hsr-9* mutants, while the average remained six, we observed ∼9% of nuclei containing only five GFP::COSA-1 foci (Figure 5A, B). These results suggest that HSR-9 regulates the processing of DSBs into COSA-1-marked events.

Given HSR-9’s enrichment on *X* chromosomes (Figure 2), we examined whether those nuclei containing five COSA-1 foci were lacking GFP::COSA-1 on the *X* chromosome pair. To that end, we labelled chromosomes with antibodies directed against the synaptonemal complex central region component SYP-1 (MacQueen et al., 2002) in worms expressing both GFP::COSA-1 and mCherry::HIM-8. Among nuclei with five GFP::COSA-1 foci, where mCherry::HIM-8 could be detected, 17.4 % (4/23) lacked a GFP::COSA-1 on the *X* chromosome pair (Figure 5C). If all six chromosome pairs have an equal probability of not receiving a COSA-1 focus, 16.7% nuclei are predicted to lack a GFP::COSA-1 foci on the *X* chromosome pair. Thus, the lack of a COSA-1 focus in nuclei containing five foci is not limited to the *X* chromosome pair in the *hsr-9* mutant.

The reduction in COSA-1 foci in *hsr-9* mutants was surprising given the high progeny viability and suggests that COSA-1 may not mark all crossovers in the absence of HSR-9. To determine whether the alteration in GFP::COSA-1 foci reflects changes in genetic crossovers, we monitored genetic linkage between SNP markers on chromosomes *I* and *V* in Bristol/Hawaiian hybrid strains (Figure 5D). There was no statistical difference between the genetic map distances in wild type and *hsr-9* for either chromosome *I* or *V* (*I*: WT = 45.1cM; *hsr-9(xoe17)* = 50.39cM; *V*: WT= 47.36; *hsr-9(xoe17)* = 40.86cM; Figure 5D, File S4), suggesting that genetic crossover numbers are not altered in the absence of HSR-9. Alternatively, genetic SNP mapping is not sensitive enough to detect subtle changes in crossover numbers distributed throughout the genome.

In *C*. *elegans*, crossovers are not evenly distributed along the length of the chromosomes but are enriched on the gene-poor arms and many meiotic mutants alter crossover distribution (Barnes et al., 1995; Lim et al., 2008; Rockman & Kruglyak, 2009). Analysis of crossover distribution revealed that mutation of *HSR-9* had little effect, except in the middle of chromosome *V* where there were statistically more crossovers in the left-center compared to wild-type hermaphrodites (2.77% vs.14.13%; p=0.0405; Table S4, Figure 5D). Together, these results suggest that HSR-9 does not alter crossover numbers but plays a role in promoting COSA-1 accumulation at a subset of recombination events.

### HSR-9 in meiotic checkpoint signaling

To determine whether HSR-9 functions in meiotic checkpoint signaling we analyzed the consequence of removing HSR-9 in mutants defective in crossover recombination leading to activation of the recombination checkpoint (Gartner & Engebrecht, 2022). To that end, we constructed *hsr-9(xoe17); zim-2(tm574)* and *hsr-9(xoe17); him-8(me4)* double mutants and analyzed progeny viability, male self-progeny, checkpoint signaling, and apoptosis. ZIM-2 binds to the chromosome *V* pairing center, while HIM-8 binds to the *X* chromosome pairing center. In *zim-2* or *him-8* mutants, chromosome *Vs* or *X* chromosomes fail to pair, synapse, and form crossovers, leading to elevated embryonic lethality or male progeny, respectively (Phillips & Dernburg, 2006; Phillips et al., 2005). Embryonic lethality was similar in *zim-2(tm574)* and the *hsr-9(xoe17); zim-2(tm574)* double mutant as well as in *him-8(me4)* and *hsr-9(xoe17); him-8(me4)* (Figure 6A). Surprisingly, we observed fewer males in *hsr-9(xoe17); him-8(me4)* compared to *him-8(me4)* (Figure 6B). A reduction in the number of males was also observed in *hsr-9(xoe17); him-8(tm611)*; *him-8(tm611)* is a deletion allele, and *hsr-9(ok759); him-8(me4)* mutants, suggesting the phenotype is not allele-specific (Figure 6B). We next examined checkpoint signaling by immunolabeling with an antibody that recognizes Ser 345 phosphorylation of the checkpoint kinase CHK-1, which is phosphorylated in response to checkpoint activation and is dependent on ATR (Jaramillo-Lambert et al., 2010). Similar levels of Ser345p, as indicated by the number of pCHK-1 foci per nucleus, were observed both in the presence and absence of HSR-9 in the *him-8* mutant background, suggesting that HSR-9 does not influence checkpoint activation in response to unpaired chromosomes and their failure to establish a crossover (Figure 6C, D). On the other hand, apoptosis was reduced in the absence of HSR-9, consistent with what was observed in response to IR (Figure 6E) (Ryu et al., 2013).

To determine the nature of the defect in production of male self-progeny, we first examined whether HSR-9 plays a role in *X* chromosome pairing by monitoring mCherry::HIM-8 in the presence and absence of HSR-9. As previously reported, in wild-type hermaphrodites, pairing of *X* chromosomes is initiated at the leptotene/zygotene (transition zone) stage of meiosis (Phillips et al., 2005). By early pachytene, stable association of HIM-8 signals was achieved in nearly 100 percent of nuclei (Figure 7A). The same pattern was observed in *hsr-9(xoe17)* mutants (Figure 7A). Thus, it is unlikely that an earlier defect in pairing is altering *X* chromosome segregation.

We next monitored oocyte chromosome nondisjunction by constructing strains containing X-linked GFP and tdTomato nuclear markers (El Mouridi et al., 2022). Hybrid strains expressing both green and red nuclear fluorescence were crossed to males carrying the *X*-linked *lon-2* mutation, allowing us to distinguish two types of non-disjunction events: oocytes containing no *X* chromosome (nullo X) fertilized by *lon-2* male sperm resulting in long males and *XX* oocytes fertilized by male sperm leading to worms expressing both nuclear GFP and tdTomato. As expected, we observed approximately equal numbers of green and red progeny from wild type and *hsr-9(xoe17)* mutants (wt: 52.5±4.4 green, 47.5±4.4 red; *hsr-9(xoe17)*: 54.4±4.0 green, 45.6±4.0 red), and recorded no nondisjunction events (Figure 7B). In *him-8(tm611)* we observed ∼38% nondisjunction events composed of 24.2±6.0% nullo *X* and 13.8±2.8 *XX* oocytes. In *hsr-9(xoe17); him-8(tm611)* we observed significantly fewer nondisjunction events (∼19%; p>0.02) of which 10.5±2.1% were nullo *X* and 8.5±3.9% were *XX* oocytes (Figure 7B). Thus, HSR-9 influences the segregation pattern of the *X* chromosomes in oocytes when their pairing, synapsis, and recombination are disrupted.

Given the role of HSR-9 in *X* chromosome segregation in the *him-8* mutant background and the enrichment of GFP::V5::HSR-9 on the *X* chromosome pair, we monitored the localization of GFP::V5::HSR-9 in the *him-8* mutant. We observed enrichment on two chromosome domains in many proliferative germ cells, as with wild type. In meiosis, many nuclei had regions with more intense fluorescence, consistent with enrichment on the unpaired *X*s, but as we could not label the *X* chromosomes (with mCherry::HIM-8 or via methods that require fixation), we could not definitively conclude these were the *X* chromosomes (Figure 7C). To gain further insight, we asked if the GFP::V5::HSR-9 enrichment was a consequence of the pairing status of the *X* chromosomes in meiosis. To that end, we examined localization of GFP::V5::HSR-9 in the *tra-2* loss-of-function mutant, which transforms *XX* animals into males (Hodgkin & Brenner, 1977). While GFP::V5::HSR-9 fluorescence was enriched on the *X*s as marked by mCherry::HIM-8 in proliferating germ cells, no enrichment was observed in meiotic cells similar to wild-type males containing a single *X* chromosome (Figure 7C). These results suggest that it is not the pairing status *per se* that leads to accumulation of GFP::V5::HSR-9 on the *X* chromosomes in oogenic germ cells.

## Discussion

We show here that HSR-9, the *C. elegans* 53BP1 homolog, is chromatin-associated, enriched on the *X* chromosome pair in oogenic germ cells, and on kinetochores at metaphase of mitosis and meiosis. Mutant analysis revealed a subtle role for HSR-9 in meiotic DSB processing, checkpoint signaling, and a previously unrecognized role in *X* chromosome segregation when X chromosomes fail to pair, synapse, and recombine.

### HSR-9 is associated with a unique chromatin state

As in mammals, HSR-9 is chromatin-associated and becomes enriched on kinetochores in dividing cells, suggesting that HSR-9 associates with a particular chromatin state. However, in contrast to mammals where 53BP1 marks DSBs in association with γ-H2AX, HSR-9 does not become enriched at either meiotic or IR-induced breaks (Figures 1 and 4A). Further, no γ-H2AX variant has been identified in *C. elegans*. In many organisms, meiotic DSBs occur at hotspots, special chromosomal sites dictated largely by the chromatin state (Tock & Henderson, 2018); however, this does not appear to be the case in *C. elegans* (Bernstein & Rockman, 2016; Kaur & Rockman, 2014). Instead, DSBs (and crossovers) are enriched on chromosome arms, which have a distinct chromatin landscape compared to the middle of the chromosomes (Lascarez-Lagunas et al., 2023). The pattern of DSBs observed may be a consequence of the holocentric nature of *C. elegans* chromosomes (Altendorfer et al., 2020). Together, these results suggest that *C. elegans* has a unique, but as yet undefined, chromatin state at DSBs sites, which does not include enrichment of HSR-9.

Although it is not enriched at DSBs, HSR-9 is enriched on *X* chromosome(s) in spermatogenic and oogenic proliferating germ cells but only the *X* chromosome pair in meiotic prophase oocytes (Figure 2). We provide evidence that HSR-9 interacts with H4K20me2/3 as 53BP1 does in mammals. However, this interaction contributes, but is not essential for, its enrichment on the *X* chromosome pair. 53BP1 has also been shown to interact with H2AK15ub (Fradet-Turcotte et al., 2013; Wilson et al., 2016), although there is no evidence that this chromatin modification is present in *C. elegans*. Thus, it is likely that HSR-9 associates with a distinct chromatin state. We do show that HSR-9, like 53BP1, has a highly disordered N-terminus, and based on results from different fixation conditions, is likely to interact dynamically with chromatin. Further, our findings that HSR-9 enrichment is specific to the *X* chromosomes in oogenic meiosis but not in male meiosis, are also consistent with its association with a unique chromatin state. That GFP::V5::HSR-9 is not enriched on the paired *X* chromosomes in male germ cells in the sex determination mutant *tra-2*, which transforms *XX* worms into males, suggests that the difference is not due to the pairing status of the *X*. This is also consistent with our previous findings that the *X* of males has distinct chromatin properties from the *X* chromosome pair in oogenesis independent of pairing status (Checchi & Engebrecht, 2011).

### The role of HSR-9 in meiotic recombination

53BP1 orthologs have been shown to function in both early and late processing of DSBs. In somatic cells, 53BP1 plays an early role in repair choice through its interaction with RIF1, the Shieldin complex, and PTIP to bind at DSBs and block resection, thereby promoting NHEJ (Chapman et al., 2013; Escribano-Diaz et al., 2013; Mirman et al., 2018; Noordermeer et al., 2018). 53BP1 has also been suggested to play later roles in both limiting RAD51 loading and stimulating strand annealing for repair by HR (Callen et al., 2020; Ferrari et al., 2020). We find that HSR-9 plays a role in meiotic DSB repair as shown by elevated levels of both RPA-1 and RAD-51 (Figure 4). This is unlikely due to a defect in repair choice, as blocking RAD-51 at DSBs by mutation of RAD-54 does not alter the number of RAD-51 foci, suggesting that the same number of DSBs are processed by HR in the *hsr-9* mutant compared to wild type. Interestingly, we find that although there appears to be a delay in the processing of DSBs in the *hsr-9* mutant, there is no effect on progeny viability suggesting that DSBs are properly repaired and establish crossovers for accurate chromosome segregation. A small subset of DSBs is processed into crossovers without accumulating COSA-1, suggesting that either COSA-1 is not required for a subset of crossovers, or more likely, that COSA-1 is required but does not accumulate to cytological visible levels in the absence of HSR-9 as suggested by Yokoo et al. (Yokoo et al., 2012). Whether this is a consequence of direct interactions between HSR-9 and DSB processing machinery, and/or an indirect effect of the chromatin state requires further investigation.

### Checkpoint signaling and sex chromosomes

In yeast, Rad9, the 53BP1 homolog, was the first checkpoint protein to be discovered and subsequent analyses of mammalian 53BP1 was consistent with a role in checkpoint signaling (Schultz et al., 2000; Weinert & Hartwell, 1989). Our study suggests that HSR-9 plays a subtle role in DNA damage checkpoint signaling. We found that in the absence of HSR-9, the checkpoint kinase, CHK-1, is phosphorylated in response to defects in chromosome pairing, synapsis, and recombination, suggesting that detection and relay through the checkpoint pathway is not perturbed. However, checkpoint-dependent apoptosis is diminished in the absence of HSR-9 as was previously shown in response to IR-induced DNA breaks (Ryu et al., 2013). Further, we did not uncover a checkpoint role for HSR-9 in response to unattached kinetochores even though HSR-9 is enriched at kinetochores. In contrast, mammalian 53BP1 has been shown to be both enriched at kinetochores and to monitor spindle attachments/tension (Jullien et al., 2002; Wang et al., 2017; Yim et al., 2017).

Although apoptosis is impaired when both autosomes and sex chromosomes are unable to pair, synapse and recombine, we discovered a reduction in male self-progeny in *hsr-9* mutants when sex chromosome pairing, synapsis and recombination are impaired. Analysis of chromosome nondisjunction events in oogenesis suggests that HSR-9 influences the pattern of *X* chromosome segregation under these conditions. Perhaps HSR-9 enrichment on the *X* chromosomes alters how the unattached *X* chromosomes align and segregate during meiosis I, leading to a higher likelihood of generating nullo *X* gametes for production of males.

## Conclusion

53BP1 regulates several aspects of chromosome biology – presumably through its chromatin association. There appears to be both commonalities and unique properties of 53BP1 homologs across evolution and these differences may reflect different chromatin environments in different organisms. In worms, its concentration on the *X* chromosomes suggests that HSR-9 plays a role in influencing *X* chromosome segregation in addition to its role in DSB processing.

## Supporting information

Strain

CRISPR

Antibodies

## Acknowledgements

We thank the Engebrecht laboratory for thoughtful discussions. We also acknowledge the Caenorhabditis Genetic Center, which is funded by the National Institutes of Health (NIH) Office of Research Infrastructure Programs (P40 OD010440) for providing strains. This work was supported by NIH GM103860 and GM103860S1 to J.E. EH was supported by the UC Davis OEOES Research Fellowship Program.

## Data availability

Strains and reagents are available upon request. The authors affirm that all data necessary for confirming the conclusions of this article are represented fully within the article and its tables and figures. Supplemental material available at Figshare. Table S1 contains *C. elegans* strain information; Table S2 contains CRISPR information; Table S3 contains antibody information and File S4 contains the raw meiotic mapping data. Figure S1 shows *hsr-9* gene structure, embryonic lethality of mutants and fusions, and localization of HSR-9::GFP::3xHA. Figure S2 shows the consequence of blocking apoptosis to *hsr-9* progeny viability, recombination analyses of the *hsr-9(ok759)*, and full gonad images of RAD-51 labelling.

---

**Figure S1.**
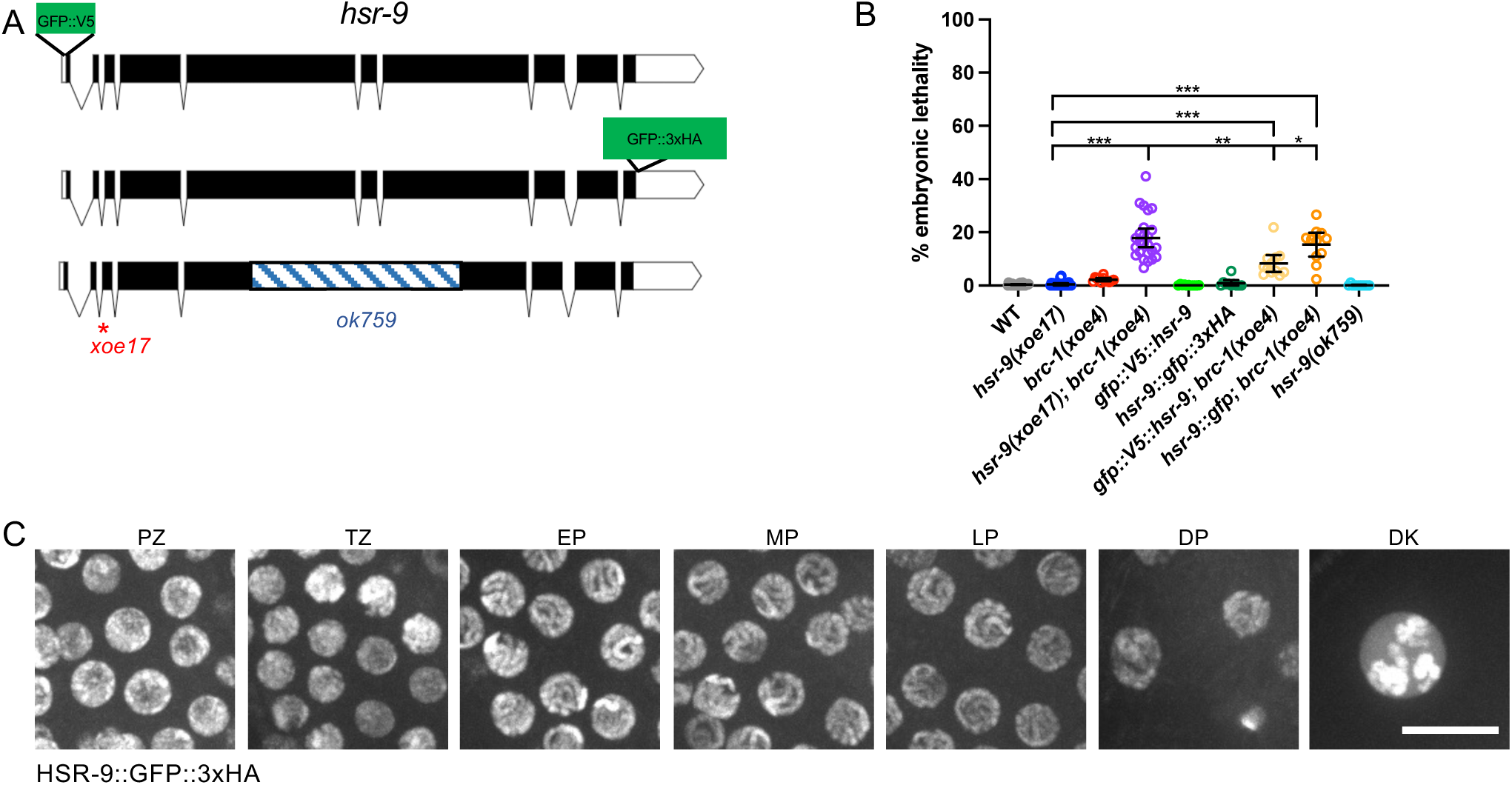
Localization and phenotype of HSR-9 fusions and mutants. A) Cartoon of *hsr-9* gene structure indicating position of fusions, and mutant alleles. B) % embryonic lethality of wild type (WT) (26), *hsr-9(xoe17)* (23), *brc-1(xoe4)* (12), *hsr-9(xoe17); brc-1(xoe4)* (25), *gfp::V5::hsr-9* (11), *hsr-9::gfp::3xHA* (10), *gfp::V5::hsr-9; brc-1(xoe4)* (12); *hsr-9::gfp::3xHA; brc-1(xoe4)* (11). Number of animals examined are in paratheses. Mean and 95% Confidence Interval shown; *** p < 0.001; ** p < 0.01; * p < 0.05 by Mann-Whitney. C) Images of HSR-9::GFP fluorescence from indicated regions of the germ line. PZ = proliferative zone; TZ = transition zone; EP = early pachytene; MP = mid pachytene; LP = late pachytene; DP = diplotene; DK = diakinesis. Scale bar = 10μm.

**Figure S2.**
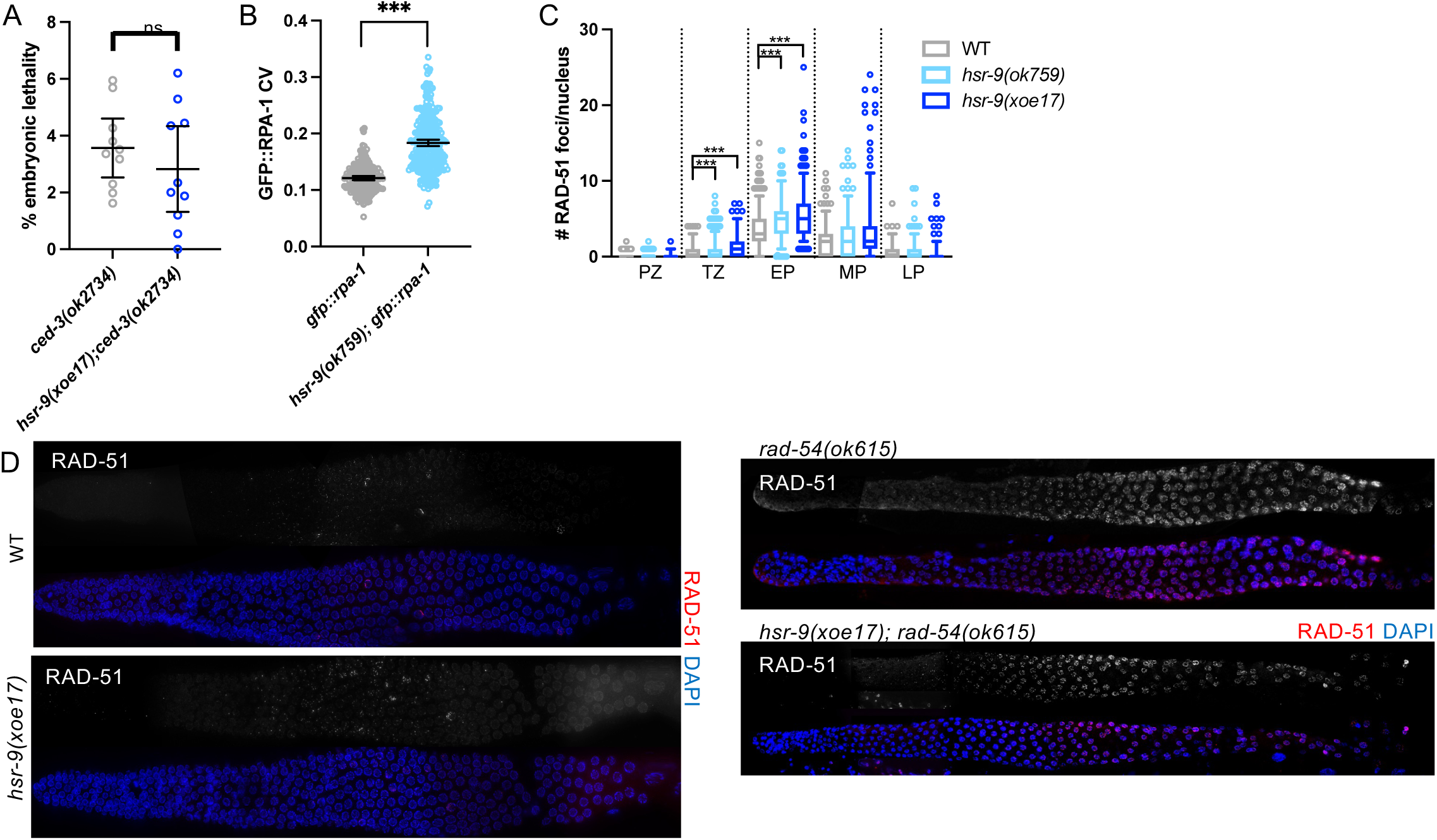
Meiotic Recombination in *hsr-9* mutants. A) Percent embryonic lethality in the apoptosis-defective *ced-3(ok2734)* mutant and *hsr-9(xoe17); ced-3(ok2734)* double mutant, n = 10 worms for each genotype. B) Coefficient of Variation (CV) of GFP::RPA-1 fluorescence was measured from 6 germlines in the pachytene region of the germ line in *gfp::rpa-1* and *hsr-9(ok759); gfp::rpa-1* worms using the 3i spinning disc microscope. C) Box whisker plots show number of RAD-51 foci per nucleus in the indicated regions. Horizontal line of each box represents the median, top and bottom of each box represents medians of upper and lower quartiles, lines extending above and below boxes indicate SD, and individual data points are outliers from 5 to 95%. Statistical significant comparisons by Mann–Whitney of WT vs. *hsr-9(ok759)* and *hsr-9(xoe17)* are indicated; ***p< 0.0001. D) Dissected germ lines immunolabelled with RAD-51 (red) and counterstained with DAPI (blue) in WT, *hsr-9(xoe17), rad-54(ok615)* and *hsr-9(xoe17); rad-54(ok615)* worms.

