## Supplementary material for "The chromatin-associated 53BP1 ortholog, HSR-9, regulates recombinational repair and *X* chromosome segregation in the *Caenorhabditis elegans* germ line": Strain

**Table S1: Strains used in this study**

| Name | Genotype | Source |
|---|---|---|
| N2 | | CGC |
| JEL1041 | *gfp::V5::hsr-9(xoe45) I* | This study |
| JEL1045 | *hsr-9::gfp::3xHA(xoe47) I* | This study |
| JEL1000 | *hsr-9(xoe17) I* | Hariri et al. 2023 |
| NB240 | *hsr-9(ok759) I* | *C. elegans* Gene Knockout Consortium |
| JEL730 | *brc-1(xoe4) III* | Li et al. 2018 |
| JEL1016 | *hsr-9(xoe17) I; brc-1(xoe4) III* | Hariri et al. 2023 |
| JEL1151 | *gfp::V5::hsr-9(xoe45) I; brc-1(xoe4) III* | This study |
| JEL1144 | *hsr-9::gfp::3xHA(xoe47) I; brc-1(xoe4) III* | This study |
| CeCL59 | *mcherry::him-8-him-8(tm611) IV* | Chenshu Li |
| JEL1327 | *gfp::V5::hsr-9(xoe45) I; mcherry::him-8-him-8(tm611) IV* | This study |
| MT14911 | *set-4(n4600) II* | Erik Andersen |
| VC997 | *set-4(ok1481) II* | *C. elegans* Reverse Genetics Facility UBC |
| JEL1322 | *gfp::V5::hsr-9(xoe45); set-4(n4600)* | This study |
| JEL1323 | *gfp::V5::hsr-9(xoe45); set-4(ok1481)* | This study |
| DH1 | *zyg-1(b1) II* | CGC |
| JEL1335 | *set-4(n4600) II; mcherry::him-8-him-8(tm611) IV* | This study |
| JEL1336 | *set-4(ok1481) II; mcherry::him-8-him-8(tm611) IV* | This study |
| RB2071 | *ced-3(ok2734) IV* | *C. elegans* Gene Knockout Consortium |
| JEL1312 | *hsr-9(xoe17) I; ced-3(ok2734) IV* | This study |
| JEL273 | *gtls2368[pie-1p::gfp::rpa-1 + unc-119(+)] II* | Sonneville *et al.,* 2012 |
| JEL1127 | *hsr-9(xoe17) I; gtls2368[pie-1::gfp::rpa-1+ unc-119(+)] II* | This study |
| JEL716 | *hsr-9(ok759) I; gtls2368[pie-1::gfp::rpa-1+ unc-119(+)] II* | This study |
| VC531 | *rad-54.L&snx-3(ok615)* I/hT2 [*bli-4(e937) let-?(q782) qls48*] (*I;III*). | CGC |
| JEL1309 | *rad-54.L&snx-3(ok615) hsr-9(xoe17)*I/hT2 [*bli-4(e937) let-?(q782) qls48*] (*I;III*). | This study |

| | | |
|---|---|---|
| JEL993 | *gfp(glo)::3xflag::cosa-1(xoe44) III* | Li *et al.*, 2022 |
| JEL1010 | *hsr-9(xoe17) I; gfp(glo)::3xflag::cosa-1(xoe44)* | This study |
| AV630 | *meIs8[unc-119(+) pie-1promoter::gfp::cosa-1] II* | Yokoo *et al.*, 2012 |
| JEL971 | *hsr-9(ok759) I; meIs8[unc-119(+) pie-1promoter::gfp::cosa-1] II* | This study |
| JEL1333 | *hsr-9(xoe17) I; gfp(glo)::3xflag::cosa-1(xoe44) III; mcherry::him-8-him-8(tm611) IV* | This study |
| CB4856 | Hawaiian | L. Hollen |
| JEL1020 | *hsr-9(xoe17) I* (Hawaiian) | This study |
| CA258 | *zim-2(tm574)* | Shohei Mitani |
| JEL1334 | *hsr-9(xoe17); zim-2(tm574)* | This study |
| | *him-8(me4)* | This study |
| JEL1341 | *hsr-9(xoe17); him-8(me4)* | This study |
| FM325 | *him-8(tm611)* | Shohei Mitani |
| JEL1348 | *hsr-9(xoe17); him-8(tm611)* | This study |
| JEL1344 | *hsr-9(ok759); him-8(me4)* | This study |
| EG7994 | *unc-119(ed3)III; oxTi395X[eft-3p::tdTomato::H2B::unc-54 3'UTR + Cbr-unc-119(+)]* | El Mouridi *et al.,* 2022 |
| EG8951 | *oxTi1015X[eft-3p::GFP::2xNLS::tbb-2 3'UTR + NeoR]* | El Mouridi *et al.*, 2022 |
| JEL1364 | *oxTi395X[eft-3p::tdTomato::H2B::unc-54 3'UTR + Cbr-unc-119(+)]/oxTi1015X[eft-3p::GFP::2xNLS::tbb-2 3'UTR + NeoR]* | This study |
| JEL1372 | *hsr-9(xoe17) I; oxTi395X[eft-3p::tdTomato::H2B::unc-54 3'UTR + Cbr-unc-119(+)]/oxTi1015X[eft-3p::GFP::2xNLS::tbb-2 3'UTR + NeoR]* | This study |
| JEL1365 | *him-8(tm611 IV); oxTi395X[eft-3p::tdTomato::H2B::unc-54 3'UTR + Cbr-unc-119(+)]/oxTi1015X[eft-3p::GFP::2xNLS::tbb-2 3'UTR + NeoR]* | This study |
| JEL1366 | *hsr-9(xoe17); him-8(tm611) IV; oxTi395X[eft-3p::tdTomato::H2B::unc-54 3'UTR + Cbr-unc-119(+)]/oxTi1015X[eft-3p::GFP::2xNLS::tbb-2 3'UTR + NeoR]* | This study |
| JEL46 | *fem-3(e19996)/nT1-GFP IV; lon-2(e678) X* | Jaramillo-Lambert and Engebrecht, 2010 |
| JEL1373 | *gfp::V5::hsr-9(xoe45) I; him-8(tm611) IV* | This study |
| CB2754 | *tra-2(e1095)/dpy-10(e128) unc-4(e120) II* | J. Hodgkin |
| JEL1363 | *gfp::V5::hsr-9 II; tra-2(e1095)/dpy-10(e128) unc-4(e120) II; mcherry::him-8-him-8(tm611) IV* | This study |

*C. elegans* Deletion Mutant Consortium, 2012 large-scale screening for targeted knockouts in the *Caenorhabditis elegans* genome. G3 11:1415-142.

El Mouridi, S., Alkhaldi, F., & Frokjaer-Jensen, C. (2022). Modular safe-harbor transgene insertion for targeted single-copy and extrachromosomal array integration in Caenorhabditis elegans. *G3 (Bethesda)*, *12*(9). https://doi.org/10.1093/g3journal/jkac184

Hariri, S., Li, Q., & Engebrecht, J. (2023). 53bp1 mutation enhances brca1 and bard1 embryonic lethality in C. elegans. *MicroPubl Biol*, *2023*.

Jaramillo-Lambert, A., & Engebrecht, J. (2010). A single unpaired and transcriptionally silenced X chromosome locally precludes checkpoint signaling in the Caenorhabditis elegans germ line. *Genetics*, *184*(3), 613-628. https://doi.org/10.1534/genetics.109.110338

Li, Q., Kaur, A., Mallory, B., Hariri, S., & Engebrecht, J. (2022) Inducible degradation of dosage compensation protein DPY-27 facilitates isolation of Caenorhabditis elegans males for molecular and biochemical analyses. G3 12:jkac085. https://doi.org/10.1093/g3journal/jkac085.

Li, Q., Saito, T. T., Martinez-Garcia, M., Deshong, A. J., Nadarajan, S., Lawrence, K. S., et al. (2018). The tumor suppressor BRCA1-BARD1 complex localizes to the synaptonemal complex and regulates recombination under meiotic dysfunction in Caenorhabditis elegans. PLoS Genet 14:e1007701 https//doi.org/10.1371/journal.pgen.1007701

Sonneville, R., Querenet, M., Craig, A., Gartner, A., & Blow, J. J. (2012)The dynamics of replication licensing in live *Caenorhabditis elegans* embryos. J Cell Biol. 196: 233-246. https://doi.org/10.1083/jcb.201110080

Yokoo, R., Zawadzki, K. A., Nabeshima, K., Drake, M., Arur, S., & Villeneuve, A. M. (2012). COSA-1 reveals robust homeostasis and separable licensing and reinforcement steps governing meiotic crossovers. *Cell*, *149*(1), 75-87. https//:doi.org/10.1016/j.cell.2012.01.052
