## Supplementary material for "The chromatin-associated 53BP1 ortholog, HSR-9, regulates recombinational repair and *X* chromosome segregation in the *Caenorhabditis elegans* germ line": CRISPR

**Table S2: CRISPR information**

<table>
<tr><td colspan="3">gfp::v5::hsr-9</td></tr>
<tr><td>Guide RNA</td><td colspan="2">ctgaactgcttggaggaaaa</td></tr>
<tr><td>Repair template</td><td colspan="2">gggcgacactcagtacatttttttgaacagcgggggcgggcgtttgaattgaattttcaaacgatttttttttgacgtcgtaactttttgtttcgcagacac<br>ttttacgcttaaaatggcaaaattaaacacatatgttacagcatagacttgaagaactgaactgcttggaggaaaatgagtaaaggagaagaact<br>ctttaccggagtcgtcccaattctcgtcgagctcgacggagacgtaaacggacacaaattctcggtttccggagaaggagaaggagatgctact<br>tatggaaaactcacccttaaattcatttgcaccaccggaaagctcccagtgccgtgggtaagtttgtgataatccaatttcaattcgcaatggtca<br>tcgtttttttcagccgacccttgtgactacattctgctacggagttcaatgcttttcccgctacccagaccacatgaagcgtcacgatttcttcaaatc<br>agccatgccagaaggatacgtccaggagcgaacaattttcttcaaggacgatggaaactacaagactcgtaagtttttactccgcttttaacaat<br>ggttgtttgacatcattttttcaggtgctgaagtcaagtttgaaggagatactctcgtgaaccgcattgagctcaagggaatcgacttcaaagaaga<br>tggaaacattcttggacacaagctcgaatacaactataactcgcacaacgtgtatatcatggccgacaagcaaaaaaatggaatcaaggtcgt<br>aagtttgatgaaacggtttcgtcttatatacactaatggtacttttcagaacttcaagattcgccacaacatcgaagacgggtcggttcaactcgct<br>gatcactaccagcagaacacaccaattggagatggaccagtcctcctccctgataatcactacctttccactcaatccgctcttagcaaggatc<br>caaatgagaaaagagatcacatggttcttctcgagtttgtcaccgccgccggaatcacccacggaatggacgagttgtacaagtccgccggag<br>gaagtggtaagcctatccctaaccctctcctcggtctagatagtactggagcatcgggagcctcaggagcatcgatggcatcgtcttcgaacact<br>atgggtaattttttcgatataataagaatgtgaaaggttaattaatatttttgcgggtcaagcatataaaattcaaccttattgaaattaaggtggtcac<br>tcgaagtgaatatcgactctaaaaattaaacttaacttacgtgaagttttccttttattttctgttttcagaattcg</td></tr>
<tr><td>Genotyping primers</td><td>Internal<br>Fwd: ACGGAAAGAAAACCCACTGC<br>Rev: GAATTGGGACGACTCCGGTA</td><td>External<br>Fwd: CCAATCTGGATCGACTGGGG<br>Rev: AGCTGGGGTCTAAGTAAGCAAA</td></tr>
<tr><td colspan="3">hsr-9::gfp::3xHA</td></tr>
<tr><td>Guide RNA</td><td colspan="2">atgatattcaatgacgcgtg</td></tr>
<tr><td>Repair template</td><td colspan="2">ggagcaagtagagtcacttccgaatgggttatccaagtgagattccgttttcctcgtcgtttaattgttatacatttttcagacaataattcttggcaa<br>agctcctgagccaaatgctcatccaaaattcgatccataccgtctgcatcatcgcacgcgtcatggagcatcgggagcctcaggagcatcgatg<br>agtaaaggagaagaactctttaccggagtcgtcccaattctcgtcgagctcgacggagacgtaaacggacacaaattctcggtttccggagaa<br>ggagaaggagatgctacttatggaaaactcacccttaaattcatttgcaccaccggaaagctcccagtgccgtgggtaagtttgtgataatccaa<br>tttcaattcgcaatggtcatcgtttttttcagccgacccttgtgactacattctgctacggagttcaatgcttttcccgctacccagaccacatgaagc<br>gtcacgatttcttcaaatcagccatgccagaaggatacgtccaggagcgaacaattttcttcaaggacgatggaaactacaagactcgtaagttt<br>ttactccgcttttaacaatggttgtttgacatcattttttcaggtgctgaagtcaagtttgaaggagatactctcgtgaaccgcattgagctcaaggga</td></tr>
</table>

<table>
<tr><td></td><td colspan="2">atcgacttcaaagaagatggaaacattcttggacacaagctcgaatacaactataactcgcacaacgtgtatatcatggccgacaagcaaaaa<br>aatggaatcaaggtcgtaagtttgatgaaacggtttcgtcttatatacactaatggtacttttcagaacttcaagattcgccacaacatcgaagacg<br>ggtcggttcaactcgctgatcactaccagcagaacacaccaattggagatggaccagtcctcctccctgataatcactacctttccactcaatc<br>cgctcttagcaaggatccaaatgagaaaagagatcacatggttcttctcgagtttgtcaccgccgccggaatcacccacggaatggacgagttg<br>tacaagtacccatacgacgtcccagactacgcctacccatatgatgtcccggattacgcttacccatacgatgttccagattacgcttgaatatc<br>atctcatctcacccccctttccccatttcccttctcattttgtacactacgtttttatatcaactttttaaataactattctactttccctaccgtgctcca<br>tccgctgaaatatcgtcgatctcttactcgaaaatttacccgc</td></tr>
<tr><td>Genotyping primers</td><td>Internal<br>Fwd: CTTGAGCTTCTCGGTGCTGA<br>Rev: GTGTCCGTTTACGTCTCCGT</td><td>External<br>Fwd: GAATGGTGGGATCGTCACGG<br>Rev: GTTGACCAGCACACTGAGCG</td></tr>
</table>
