## Supplementary material for "The chromatin-associated 53BP1 ortholog, HSR-9, regulates recombinational repair and *X* chromosome segregation in the *Caenorhabditis elegans* germ line": Antibodies

**Table S3. Antibody information**

| Target | Host | Dilution | RRID | Source |
|---|---|---|---|---|
| GFP | Rabbit | 1:500 | AB_303395 | Abcam/ab290 |
| H4K20me1 | Rabbit | 1:250 | AB_306967 | Abcam/ab9051 |
| HCP-3 | Rabbit | 1:500 | AB_1235828 | SDIX/2954.00.02 |
| phospho-Histone H3 (Ser10) | Rabbit | 1:200 | AB_310177 | Sigma-Aldrich/06-570 |
| RAD-51 | Rabbit | 1:5,000 | AB_2616441 | SDIX/2948.00.02 |
| SYP-1 | Rabbit | 1:200 | | Villeneuve lab |
| pCHK-1 | Rabbit | 1:50 | AB_2080472 | Santa Cruz/sc-17922 |
